# Automated model building and protein identification in cryo-EM maps

**DOI:** 10.1101/2023.05.16.541002

**Authors:** Kiarash Jamali, Lukas Käll, Rui Zhang, Alan Brown, Dari Kimanius, Sjors H.W. Scheres

## Abstract

Interpreting electron cryo-microscopy (cryo-EM) maps with atomic models requires high levels of expertise and labour-intensive manual intervention. We present ModelAngelo, a machine-learning approach for automated atomic model building in cryo-EM maps. By combining information from the cryo-EM map with information from protein sequence and structure in a single graph neural network, ModelAngelo builds atomic models for proteins that are of similar quality as those generated by human experts. For nucleotides, ModelAngelo builds backbones with similar accuracy as humans. By using its predicted amino acid probabilities for each residue in hidden Markov model sequence searches, ModelAngelo outperforms human experts in the identification of proteins with unknown sequences. ModelAngelo will thus remove bottlenecks and increase objectivity in cryo-EM structure determination.

## Introduction

Knowledge of the three-dimensional atomic structures of proteins and nucleic acids is pivotal for our understanding of the molecular processes of life. In recent years, considerable advances have been made in the determination of structures of biological macromolecules by electron cryo-microscopy (cryo-EM), culminating in cryo-EM maps of proteins with sufficient resolution to resolve individual atoms (*1, 2*). Accordingly, the number of new cryo-EM structures in the electron microscopy database (EMDB) (*3*) is growing exponentially. If this trend continues, approximately 100,000 cryo-EM structures will be determined in the next 5 years (*4*).

Over two-thirds of last year’s (2022) structures had resolutions better than 4 Å. Although individual atoms are not resolved at resolutions between 2-4 Å, reliable atomic models can be built by exploiting prior knowledge of the chemical structures of the proteins and nucleic acids in the sample, including their amino acid and nucleic acid sequences. Typically, atomic model building in cryo-EM maps is performed using manual procedures in three-dimensional computer graphics programs (*5, 6*). Atomic model building is often time-consuming and requires substantial levels of expertise to produce accurate models. At resolutions better than 3 Å, experts can build atomic models with few errors, whereas at resolutions below 4 Å, avoiding mistakes is challenging. It is therefore not uncommon for atomic models of biological complexes to contain errors (*7*), with potentially grievous consequences (*8*).

Structure determination by cryo-EM is also an increasingly important tool for the discovery of new subunits in biological complexes. Because of its relaxed requirements for sample quantity and purity compared to other structural biology techniques, cryo-EM is capable of determining structures of complexes purified from endogenous sources. Many such complexes contain subunits of unknown identities. Without prior knowledge of the amino acid sequence, identifying the chemical identity of individual amino acids in cryo-EM maps is difficult, and requires relatively high resolutions. Yet, provided one can build stretches of several consecutive amino acids, database searches with the sequence fragments can lead to the identification of the corresponding protein. Recent examples include the identification of TMEM106B in amyloid filaments from human brains (*9–11*) and the detection of subunits of axonemal complexes (*12, 13*).

Here, we introduce a machine-learning approach, called ModelAngelo, for the automated building of atomic models and the identification of proteins in cryo-EM maps. Machine learning approaches often require large amounts of training data. For example, recent protein language models were trained on tens of millions of sequences (*14*) and AlphaFold2 was trained on more than 200,000 structures (*15*). In contrast, fewer than 13,000 cryo-EM structures with resolutions better than 4 Å have been determined to date and many of these are redundant. The limited amount of available training data prompted us to design a multi-modal machine-learning approach that combines local information from the cryo-EM map surrounding each protein or nucleic acid residue with additional information from the protein sequences in the sample and the local geometry of the structure. Similar sources of information are used by human experts when manually building atomic models in cryo-EM maps.

The sudden availability of atomic models for millions of proteins from protein structure prediction by AlphaFold2 (*15, 16*) has helped to guide and accelerate model building (*17*). However, previous attempts to fully automate atomic modelling (*18–24*) or the identification of unknown proteins (*25–27*) have so far failed to become mainstream, although DeepTracer (*21, 24*) and findMySequence (*25*) have gained some traction. Still, atomic modelling remains a time-consuming and expert-dependent process in many structure determination projects. With the ongoing exponential growth in cryo-EM structures and the continuing influx of newcomers to the cryo-EM field, automation will be key in removing bottlenecks and replacing the dependence on human experts with objective methods that are accessible to all. In what follows, we demonstrate that ModelAngelo can meet this need. Although subsequent error checking and refinement remain necessary, ModelAngelo outperforms human experts in identifying unknown proteins and produces initial atomic models of comparable completeness to those obtained by human experts.

## Approach

Automated model building of proteins and nucleic acids in ModelAngelo comprises three steps (Figure 1a). Details about the network architectures that underlie these steps and how they are trained have been described in (*28*).

**Figure 1:**
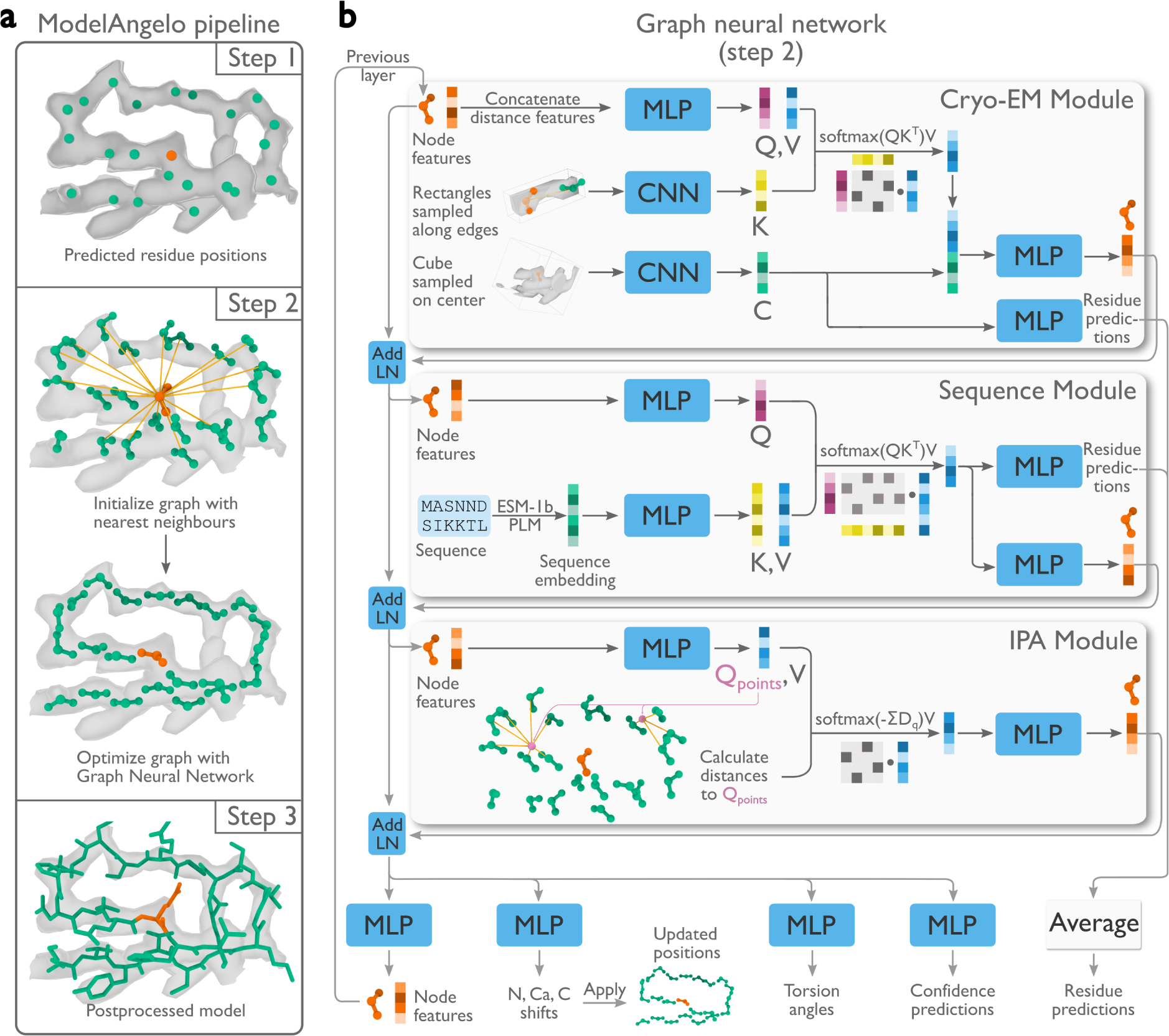
**Atomic modelling in ModelAngelo. a**, Three steps build atomic models: 1) A convolutional neural network (CNN) predicts protein and nucleic acid residue positions; 2) A graph neural network (GNN) optimises these positions and orientations (panel b); 3) A postprocessed, optimised graph forms a complete atomic model. **b**, The GNN, arranged in eight layers with three modules, uses a feature vector per residue passed through multi-layer perceptrons (MLP), integrated with additional data via attention mechanisms that have query (Q), key (K), and value (V) vectors. Stable gradient propagation is ensured by residual connections with layer norms (Add LN) (*29*). Residue feature vectors are used to update residue positions and orientations. They are also used to predict torsion angles, confidence scores, and residue identities at the end of each layer.

### Graph initialisation

Positions for the backbone C*α* atom of amino acids and the phosphor atom of nucleic acids are predicted using a convolutional neural network (CNN). This CNN is a modified feature-pyramid network (*30*) that predicts whether each voxel in the cryo-EM map contains the C*α* atom of an amino acid, the phosphor atom of a nucleic acid residue, or neither. A graph is then constructed, where each residue is a node, and edges are formed between each residue and its twenty nearest neighbours.

### Graph optimisation

A graph neural network (GNN) is used to optimise the positions and orientations of the residues, to predict their amino or nucleic acid identity, and to predict torsion angles for their side chains or bases. The GNN consists of three modules: a cryo-EM module, a sequence module, and an invariant point attention (IPA) module (Figure 1b). Each node of the graph is associated with a residue feature vector. Each module takes the residue feature vector as input, combines it with new information, and outputs an updated residue feature vector that is passed to the next module. The sequential application of the three modules in eight layers (Figure 1b) allows the gradual extraction of more information from the different inputs.

The cryo-EM module incorporates information from the cryo-EM map and comprises two parts. Firstly, the input feature vector is passed through a multi-layer perceptron (MLP) network to generate query and value vectors. These vectors are used for cross-attention (*31*) with key vectors that are calculated from a CNN on rectangular boxes that are extracted from the cryoEM density map that point from the current residue to its twenty nearest neighbours. Intuitively, the cross-attention mechanism allows mixing information from each residue with that of its twenty nearest neighbours, depending on whether the cryo-EM density between them looks connected. Secondly, a cubic box is extracted from the cryo-EM map around the position of the current residue and passed through another CNN. The resulting vector is used in two ways: to generate amino and nucleic acid identity predictions through an MLP; and after concatenation with the vector from the cross-attention, it is passed through another MLP to generate the output residue feature vector of the cryo-EM module.

The sequence module performs cross-attention for each residue with the user-provided amino acid sequences, which are embedded using the pre-trained protein language model, ESM1b (*32*). This incorporates information that is learned by the language model from many amino acid sequences, including multiple homologues. The information in protein language models has been shown to be sufficient for protein structure prediction (*14*). The vector from the crossattention is used in two ways: a first MLP is used to generate amino and nucleic acid identity predictions; a second MLP generates the output residue feature vector of the sequence module. For nucleic acid residues, the sequence module is not used.

The IPA module incorporates information from the geometry of the nodes in the graph and was inspired by the module with the same name in AlphaFold2 (*15*). An MLP calculates four query points per residue and the Euclidean distance between the query points and the location of the neighbouring nodes is used to replace the cosine similarity of the attention algorithm between the query and key vectors. Intuitively, this allows the model to learn information about the topology of neighbouring residues, for example about secondary structure. In fact, disabling this module in an ablation study led to atomic models with incorrect secondary structure geometry (*28*).

### Post-processing

The residue feature vectors are post-processed to generate an atomic model. The feature vectors are used as inputs into two separate MLPs to predict new positions and orientations for each residue, as well as torsion angles for amino acid side chains and nucleic acid bases. They are also used to predict a confidence score for each residue, which is based on the network’s predicted root-mean-square deviation (RMSD) for the backbone atoms with the deposited structure. In addition, the predictions for the amino or nucleic acid identities from the cryo-EM and sequence modules are averaged to generate probabilities for each possible identity for all residues. These vectors are converted into a hidden Markov model (HMM) profile (see below) that is used for a search against the input sequences using HMMER (*33*). Matched residues, as defined in (*34*), are mutated to the corresponding amino or nucleic acid in the input sequences, and separate chains are connected based on their assigned sequences and proximity. Finally, chains shorter than four residues are pruned from the model, and a full atomic model is generated from the predicted positions and orientations of each residue and their corresponding amino acid or nucleic base torsion angle predictions, using idealised geometries. The predicted backbone RMSD values are mapped to a score between 0 and 1, corresponding to a linear range for RMSDs between 1.2 and 0.5 Å, respectively. This score is stored in the B-factor column of the output coordinate file as a measure of local confidence in the backbone geometry.

### Generating HMM profiles

A profile HMM is a probabilistic model representing the multiple sequence alignment (MSA) of a set of related sequences. The parameters of a profile HMM are normally estimated from the MSA it strives to model, however, here, they are instead estimated from ModelAngelo predictions. There are three types of states in the profile HMM. For each position of the MSA’s consensus sequence, there is a match (*M*), a delete (*D*), and an insert (*I*) state with respect to the query sequences. There are two types of probabilities in a profile HMM: transition and emission. The transition probabilities reflect the probability of a sequence going between the *M*, *I*, and *D* states from one position of the profile to the next. ModelAngelo uses the confidence metric, *c*^(^*^i^*^)^, that it predicts for each residue *i* to construct the transition probabilities as follows:

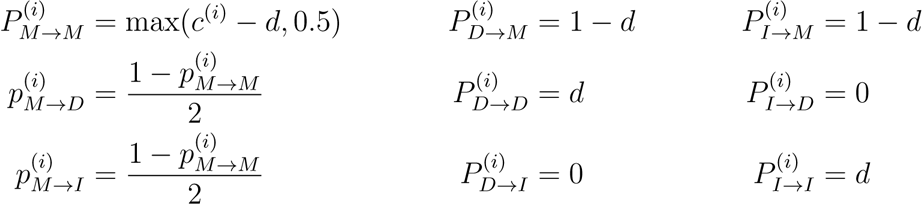

The strategy to set 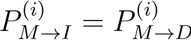; the constant *d* = 0.5; and the minimum value of 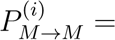 0.5 were chosen arbitrarily and are never optimised. The emission probabilities represent the probability of each amino acid being produced in a *M* or *I* state. For these, ModelAngelo uses its predicted probability distribution of the amino acids for each residue. The resulting HMM profiles are compatible with HMMER3 (*35*) and HHblits (*36*).

### Recycling

Inspired by AlphaFold (*15*), we recycle the post-processed model from one round of the GNN as the starting point of a subsequent round of graph optimisation. For this purpose, ModelAngelo was trained with a random number of 1-3 recycling steps. During inference, we perform three rounds of recycling, as the performance plateaus after three rounds.

### Training

ModelAngelo was trained on maps deposited in the EMDB (*3*) before April 1st, 2022 with resolutions better than 4 Å and paired with models in the PDB (*37*) that cover the entire map correctly, as detailed in (*28*). PDB files that included insertion codes, i.e. additional residues relative to the reference sequence, were removed. This resulted in 3715 map-model pairs that were used during training. All cryo-EM maps were resampled to a common pixel size of 1 Å . For comparison, findMySequence only uses 117 pairs, while DeepTracer uses approximately 1400 (*21, 25*).

### Protein identification

To allow model building for structures with unknown sequences, we also trained a version of ModelAngelo without its sequence module. Still, for each protein residue, ModelAngelo predicts probabilities for all twenty amino acids. Within ModelAngelo, these probabilities are converted into HMM profiles and used for searches in HMMER3 (*35*) as described above, but using a larger proteome, rather than only the sequences known to be present in the structure.

## Results

### ModelAngelo builds protein models of comparable quality to those built by humans

To test ModelAngelo, we first considered all cryo-EM structures determined to at least 4 Åresolution and released from the EMDB between the cutoff date for training, April 1st, 2022, and February 9th, 2023. To reduce computational costs, we excluded structures with more than 30,000 protein residues. We also removed viruses with icosahedral symmetry, for which typically only the asymmetric unit was built. To ensure none of the sequences were seen before during training, we removed structures that had protein chains with more than 10% sequence identity to any of the proteins in the training set. Finally, we removed structures with insertion codes and other irregularities. This resulted in a test set of 177 structures, on which we ran ModelAngelo. Using a single A100 GPU, the smallest structure (PDB ID: 8DWI, with a molecular weight of 54.7 kDa) took 2 minutes; the largest structure (PDB ID: 7UMS, with a molecular weight of 1.85 MDa) took 53 minutes. The output coordinates from ModelAngelo were refined against the cryo-EM map using a standard refinement cycle in Servalcat (*38*), and the refined models were compared to the deposited ones.

To assess the quality of the models generated by ModelAngelo, we analyzed the Q-scores (*39*) of all structures in the test set. The Q-score measures the resolvability of individual atoms in cryo-EM maps, thus reflecting the quality of the built model. Provided the model is built well, Q-scores also correlate with the local resolution, which can vary in cryo-EM maps: Q-scores of 0.4 are typical for cryo-EM maps at 4 Å resolution, values of 0.6 for maps at 3 Å, and values better than 0.7 for maps beyond 2 Å resolution (*39*). We implemented Q-score calculation in ModelAngelo and calculated average Q-scores for all atoms in each residue of both the deposited models and those built by ModelAngelo. Next, we calculated backbone root mean squared deviations (RMSDs) between the protein models built by ModelAngelo and those deposited, and plotted these against the Q-scores of the deposited residues (Figure 2a; pink line). As expected, ModelAngelo builds models with lower RMSDs for residues with higher (better)Q-scores. Even for residues with Q-scores as low as 0.4, ModelAngelo builds models with backbone RMSDs lower than 1.0 Å . We also measured the completeness of the models built by ModelAngelo. We define completeness as the fraction of residues that are built with their C*α* atom within 3 Å of the deposited model and with the correct amino acid assignment. As with backbone RMSD, completeness improves for residues with higher Q-scores (Figure 2a; blue line). Overall, ModelAngelo built 77% of all 410,585 residues in the test set. Analysis of the deposited Q-scores shows that those residues not built by ModelAngelo have lower Q-scores than those that are built (Figure 2b). In the deposited models, many of the residues with the lowest Q-scores were probably obtained by rigid-body docking of protein domains into poorly resolved regions of the cryo-EM maps. Excluding the 51,446 residues with Q-scores below 0.4, ModelAngelo built 85% of the residues in the test set. A comparison of Q-scores calculated for the models built by ModelAngelo with those calculated for the deposited models shows that models from ModelAngelo are of similar quality to the deposited ones (Figure 2c). The same is also true for overall Fourier shell correlation values between the cryo-EM maps and those parts of the models that were both built by ModelAngelo and present in the deposited models (Figure 2d).

**Figure 2:**
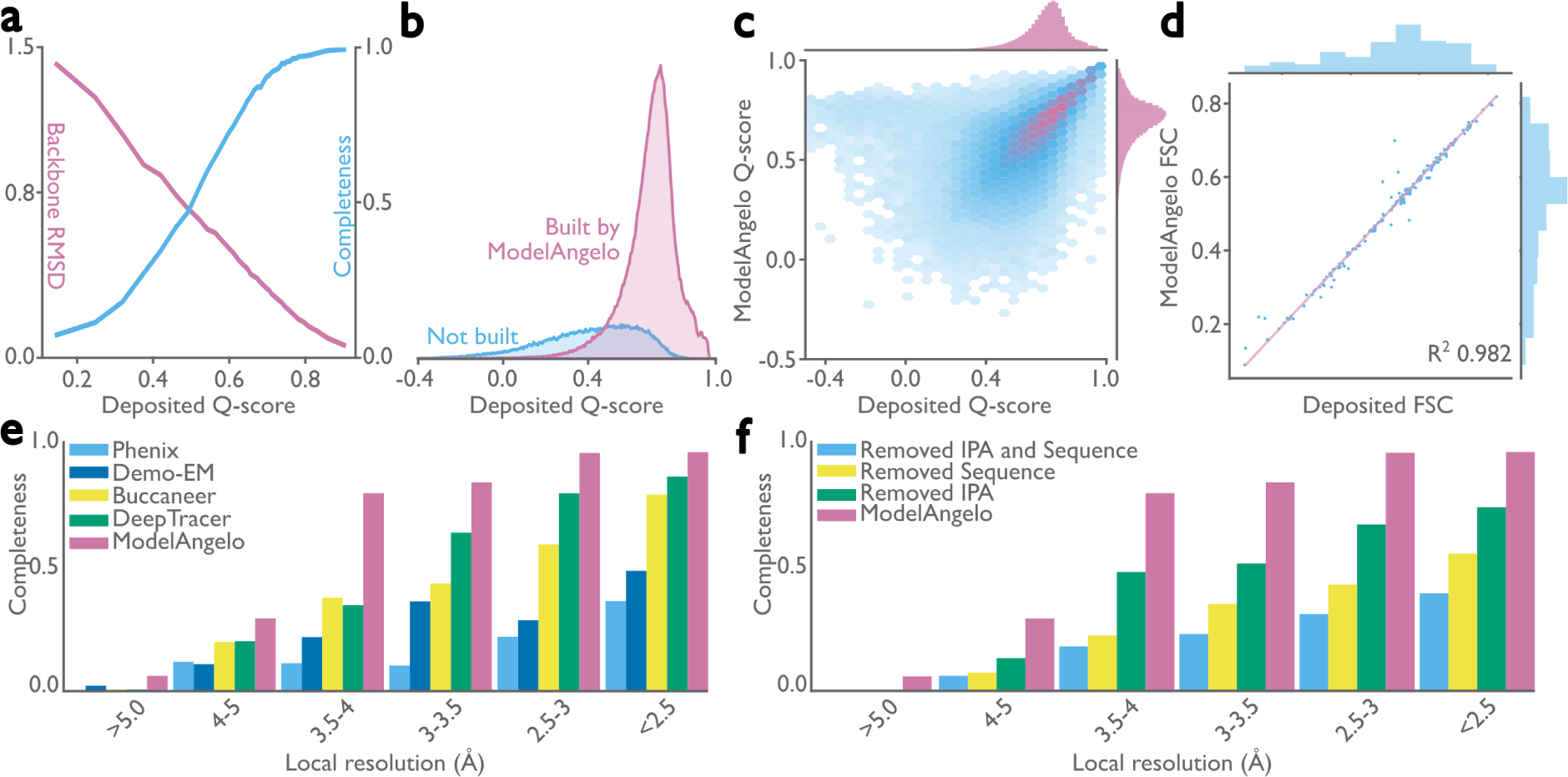
**Performance of ModelAngelo for atomic modelling of proteins. a**, Backbone root-mean-square deviation (RMSD) and model completeness plotted as a function of the target model Q-scores. **b**, Q-score distribution of residues in the deposited models, comparing those built by ModelAngelo with those not built. **c**, Q-score comparison between ModelAngelo predicted models and the deposited models. **d**, Model-to-map Fourier shell correlation (FSC), as calculated by Servalcat (*38*), after refining both models and using only residues present in both ModelAngelo and deposited models. **e**, Model completeness for various automated modelbuilding softwares for different local resolution ranges in the maps. **f**, Model completeness for ModelAngelo and versions of ModelAngelo where its Sequence and/or IPA modules were ablated. Panels a-d relate to the test set of 177 structures; panels e and f to the subset of 27 structures.

### ModelAngelo outperforms alternative approaches

In a second test, we compared the performance of ModelAngelo with existing approaches for automated model building in cryo-EM maps. For this test, we used a subset of 27 protein structures from the 177 structures described above. We selected nine single-chain structures, nine homo-oligomeric structures, and nine hetero-oligomeric structures. For each of these types of structures, we selected three structures with overall resolutions below 3.3 Å, three structures between 3.3 and 2.8 Å, and three structures with resolutions better than 2.8 Å . For all 27 structures, unfiltered half-maps were available for download from the EMDB, and we used these to calculate local resolutions in ResMap (*40*). We then used Phenix (*41*), Demo-EM (*42*), Buccaneer (*20*), and DeepTracer (*21*) for automated model building in these maps and compared the completeness of the resulting models with those obtained using ModelAngelo (Figure 2e and Extended Data Table 1). The best alternative approach, DeepTracer, built approximately 80% of the deposited residues in regions of the maps with local resolutions in the range of 2.5-3 Å ; the remaining approaches built models with considerably lower completeness. In contrast, ModelAngelo built up to 80% of the deposited residues in regions of the maps with local resolutions down to 3.5-4 Å, reflecting the observation that manual building by human experts also becomes prone to errors at resolutions below 4 Å . Tests, where we ran ModelAngelo without one or more of its modules, indicate that its performance comes from a combination of all three modules (Figure 2f), which is in accordance with previous observations (*28*).

### ModelAngelo builds good nucleic acid backbones

The test set of 177 structures described above contained only 103 nucleic acid chains, many with just a few nucleotides. Therefore, instead of conducting a systematic analysis as done for the proteins, we present a few test cases to illustrate the quality of nucleotide building (Figure 3). We applied ModelAngelo to 11 different ribosome structures that were determined to resolutions ranging from 1.98 to 3.80 Å (Figures 3a,b), as well as a CRISPR-associated transpososome from *Scytonema hofmanni* (Figures 3c,d) (*44*). Although ribosome structures were included in ModelAngelo’s training set, the nucleotide sequences were not. When plotting backbone RMSDs and backbone completeness against the Q-scores of the deposited nucleotide coordinates (Figure 3e), we observed similar trends as for the protein chains. Backbone RMSDs range from 2 Å in the worst regions of the map to values better than 0.5 Å in the best regions. Likewise, near-complete backbones are built in the best regions, while backbone completeness drops to below 80 % for the worst regions. However, ModelAngelo struggles to distinguish between the two purines or the two pyrimidines, echoing the difficulty humans face in building nucleotide sequences based solely on the cryo-EM density, if the resolution does not extend beyond 2.5 Å . Consequently, when considering only correctly built sequences, the completeness of the models built by ModelAngelo drops to 80 % for the best parts of the map, and to as low as 20 % for the worst parts (Fig 3E). Users should therefore carefully validate the nucleotide chains of models built by ModelAngelo, for example by using nucleotide secondary structure predictors (*45–49*). Nonetheless, ModelAngelo considerably accelerates the process of building the nucleotide backbone, as subsequent nucleotide base changes can be made with minimal manual intervention. For the CRISPR-associated transpososome and 3 of the 11 ribosomes described above, we also used DeepTracer (*26*) and CryoREAD (*50*). ModelAngelo produced nucleotide models that were more complete and more accurate than these alternative approaches (Extended Data Table 2).

**Figure 3:**
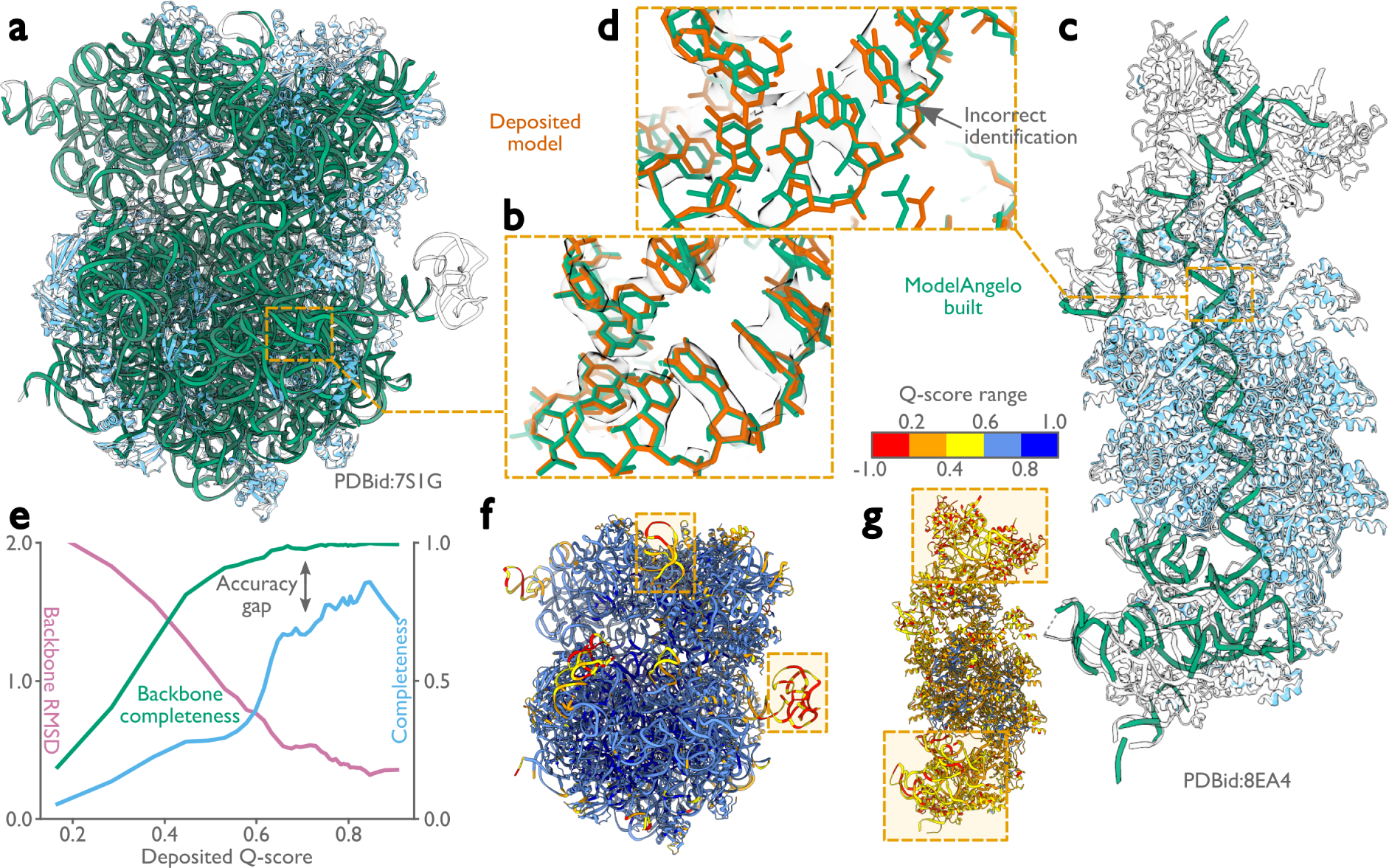
**Performance of ModelAngelo for atomic modelling of nucleic acids. a**, *Escherichia coli* ribosome built by ModelAngelo (with ribosomal RNA in green and proteins in blue) compared with the deposited model (PDB ID: 7S1G, black outline) (*43*). **b**, Zoomed-in view with nucleotide bases showing high accuracy compared to the deposited model (orange). **c**, ModelAngelo model of the V-K CAST transpososome from *Scytonema hofmanni* compared with the deposited model (PDB ID: 8EA4) (*44*). Sections not built by ModelAngelo (black outline) are in regions of low Q-score (see panel g). **d**, Zoomed-in view comparing the nucleotide bases of both models showing a sequence incorrectly identified by ModelAngelo. **e**, Backbone RMSD, backbone completeness, and sequence completeness plotted against the deposited Qscore for six ribosome structures. **f, g,** Deposited models for the structures in a and c, coloured by Q-score, with low Q-score regions boxed.

### ModelAngelo identifies protein chains that were not built by human experts

To illustrate the performance of ModelAngelo in identifying protein chains in cryo-EM maps, we applied ModelAngelo to two examples of large cryo-EM structures that were recently determined from endogenous sources. The first example is a structure of the supercomplex of the phycobilisome (PBS), photosystem I and II (PSI and PSII), and the transmembrane light-harvesting complexes (LHCs) that was imaged *in situ* in the red alga *Porphyridium purpureum* (*51*). The second example is a structure of the ciliary central apparatus and radial spokes of the green alga *Chlamydomonas reinhardtii* that was obtained by single-particle analysis following purification from cilia (*12, 13*).

At 16.7 MDa, the PBS-PSII-PSI-LHC supercomplex is one of the largest complexes determined by single-particle cryo-EM. The deposited model (PDB ID: 7Y5E) consists of 158,730 residues in 81 unique protein chains, including six chains for which the authors were unable to identify the corresponding protein. The unidentified chains were termed L_PP_1 (linker of PBS–PSII 1); CNT (for “connector”); PsbW and Psb34 (two of the core subunits of PSII); LRH (a linker protein); and L_PS_1 (photosystem linker protein 1). To identify these chains, we ran ModelAngelo without using its sequence module (using the `build no seq` option) to calculate an initial atomic model with HMM profiles for all chains, and we searched these profiles against the proteome constructed in (*52*) (using the `hmm search` option). Because of local pseudo-symmetry, all six unidentified proteins occur more than once in the cryo-EM map. This allows us to bootstrap weaker individual hits by cross-referencing their matches to the other instances. Specifically, the same six protein chains were identified for all instances, with E-values in the range of 5.8e-66 to 6.4e-2. Using the backbone traces in the deposited model, findMySequence (*25*) identified only two of the unassigned proteins (Psb34 and PsbW). Using the backbone traces generated by ModelAngelo, it also found LRH. We then constructed an input sequence file that included all chains in the deposited model plus the six newly identified chains and ran ModelAngelo again. This calculation took 23 hours on an A100 GPU. The resulting model, containing 110,742 residues, is shown in Figure 4a. For most sections of the unidentified chains, ModelAngelo built better models than those in the deposited structure, most notably for LRH and CNT. ModelAngelo did not build models for parts of the unidentified proteins that were in regions of poor cryo-EM density. Besides excellent agreement between side chain densities in the cryo-EM map and the predicted sequences (Extended Data Figure 1), the structures built by ModelAngelo were also highly similar to AlphaFold2 predictions for the unidentified chains (*15, 53*) (Extended Data Figure 2). ModelAngelo did not attempt to build amino acid or nucleotide residues in the densities for phycocyanobilin or phycoerythrobilin cofactors (Extended Data Figure 3). Because the cryo-EM maps ModelAngelo was trained on did contain cofactor densities, but it was trained to build protein and nucleic acid residues, ModelAngelo has been incentivized to ignore cofactor densities.

**Figure 4:**
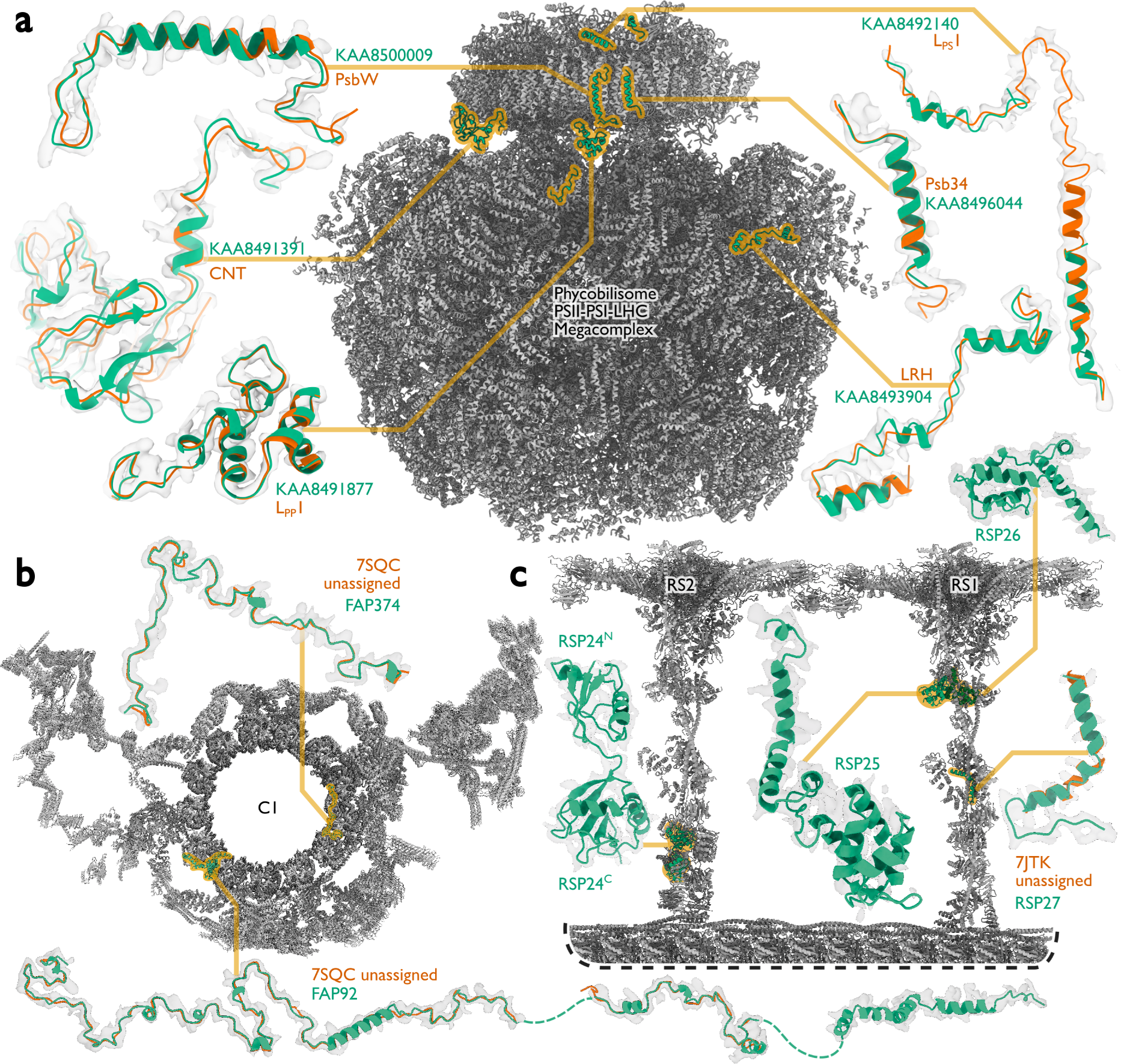
**Examples of protein identification by ModelAngelo. a**, The ModelAngelo model of the single-PBS-PSII-PSI-LHC’s supercomplex (gray) showing the positions, models, and map densities of six newly identified proteins (green). Backbone traces in the deposited model (PDB ID: 7Y5E) are shown in orange. **b**, Atomic model of the central apparatus microtubule C1 showing the positions, models, and map densities of two newly identified proteins: FAP92 and FAP374. Orange cartoons represent poly(UNK) chains deposited in the original model (PDB ID: 7SQC). **c**, An atomic model of radial spokes 1 and 2 (RS1 and RS2) bound to a doublet microtubule (gray) showing the positions, models, and map densities of four proteins (RSP24-27, green) newly identified by ModelAngelo. Only RSP27 had a backbone trace in the deposited model (orange).

Like the PBS-PSII-PSI-LHC supercomplex, the central apparatus (CA) and radial spoke complexes isolated from *C. reinhardtii* ciliary axonemes are large complexes with poorly characterized subunit compositions. Although recent cryo-EM structures had identified 23 different radial spoke proteins (RSPs) and 48 different CA proteins (*12, 13*), the deposited maps (EMD22475, EMD-24481, and EMD-25381) contained densities that were left unassigned despite considerable manual effort. To identify these proteins, we applied ModelAngelo without using its sequence module to the deposited maps and searched the resulting HMM profiles against the latest version of the *C. reinhardtii* predicted proteome (*54*) (Figure 4b and Supplementary Information). This approach identified four additional radial spoke proteins: FAP109, Cre05.g240450, Cre08.g800895, Cre17.g802036), which we rename to RSP24, RSP25, RSP26 and RSP27, respectively, and two additional CA proteins (FAP92 and FAP374) (Extended Data Table 3). Using ModelAngelo’s backbone traces, findMySequence (*25*) was unable to identify any of these proteins. Neither RSP24 (Cre08.g800895) nor RSP26 (Cre17.g802036) were annotated in earlier versions of the *C. reinhardtii* genome, explaining their absence from proteomic studies, and demonstrating the importance of high-quality genome annotations for *de novo* identification of proteins by cryo-EM. RSP27 (Cre05.g240450) was identified from a fragment of just 33 residues, demonstrating the power of ModelAngelo to identify proteins from small sections of well-resolved density. Both CA proteins (FAP92 and FAP374) bind directly to the microtubule surface and have tertiary structures poorly predicted by AlphaFold2 (Extended Data Figure 4); sidechain density was, therefore, essential for their successful identification (Extended Data Figure 5). The identification of these proteins will allow their functional relevance to the regulation of ciliary motility to be investigated through targeted genetic manipulation.

## Discussion

ModelAngelo automates atomic modelling in cryo-EM maps, building protein models of comparable quality to those built by human experts and nucleic acid models with near-complete and accurate backbones. ModelAngelo outperforms existing approaches for the automated modelling of both proteins and nucleotides. Furthermore, ModelAngelo builds these models within hours on a modern GPU, thereby removing an important bottleneck in cryo-EM structure determination. Future incorporation of ModelAngelo into automated cryo-EM image processing pipelines (*55–61*), will enable users to go from data acquisition to atomic models in a single automated procedure.

By introducing objectivity in the model-building process, ModelAngelo also informs which parts of the map can be confidently interpreted with an atomic model and which should be left uninterpreted. In this way, ModelAngelo will not only reduce the number of errors in atomic models but also play a role in making cryo-EM structure determination more accessible to the large numbers of newcomers that the field has experienced in recent years. Still, some degree of human supervision and intervention will remain necessary. Models from ModelAngelo will still need refinement, for example in Servalcat (*38*) or Phenix (*41*), to optimise their stereochemistry and fit to the cryo-EM map. Users are also strongly encouraged to manually check the output of ModelAngelo, particularly for those parts of cryo-EM maps with resolutions worse than 3.5-4.0 Å, and rigid body fitting of known domains or connecting loops in lower-resolution map regions to obtain a more complete model falls outside the scope of ModelAngelo. Colouring the model by its predicted confidence in backbone geometry, as stored in the B-factor column of the coordinate file, may guide the user towards parts of the model that are less reliable. ModelAngelo was trained with augmentation through a variety of positive and negative Bfactors. It should therefore be relatively stable to local variations in B-factor. It is possible that combining ModelAngelo with neural networks that make cryo-EM maps look more like proteins, e.g. (*62,63*), could lead to further improvements, although this would probably require re-training of ModelAngelo to reach its full potential.

Besides accelerating cryo-EM structure determination and providing objectivity in atomic modelling, ModelAngelo also identifies protein chains in cryo-EM maps better than human experts. The reason why ModelAngelo outperforms the human expert in this task likely lies in the implementation of its sequence searches. While human experts typically base their identifications on discrete assignments of individual amino acids to various residues in unknown chains, ModelAngelo exploits predicted probabilities for all twenty amino acids for every protein residue and combines this information with its predicted confidence in each residue in a full HMM search. This not only allows better identification of unknown chains but also helps ModelAngelo during the building of atomic models with known sequences, where it may potentially outperform human experts in placing protein chains for which ambiguity exists, for example when multiple homologous chains coexist in a single structure. The ability to identify proteins in cryo-EM maps will increase in importance as ongoing advances in sample preparation, microscopy, and image processing allow ever more structures to be determined for samples purified from native sources or visualised in situ by electron tomography of frozen cells or thin tissue sections.

## Acknowledgements

We thank George Ghanim, Joe Greener, Katerina Naydenova, Johannes Schwab, Zala Sekne, Sofia Lövestam, and Keitaro Yamashita for helpful discussions; Miao Gui for contributions to atomic modelling of the ciliary axonemes; and Jake Grimmett, Toby Darling, and Ivan Clayson for help with high-performance computing. This work was supported by the Medical Research Council as part of the United Kingdom Research and Innovation (MC UP A025 1013 to SHWS); the European Union’s Horizon 2020 research and innovation programme (under grant agreement no. 895412 to DK); the National Institutes of Health (R01-GM141109 to AB and R01-GM138854 to RZ); and the Knut and Alice Wallenberg Foundation (2022.0032 to LK). For the purpose of open access, the MRC Laboratory of Molecular Biology has applied a CC BY public copyright licence to any Author Accepted Manuscript version arising.

## Author contributions

KJ designed and implemented ModelAngelo. LK designed and implemented the HMM search algorithm. RZ and AB analyzed ciliary axoneme data. DK and SHWS jointly supervised the project. All authors contributed to the writing of the manuscript.

## Code availability

ModelAngelo is freely available under the open-source MIT license and can be downloaded from https://github.com/3dem/model-angelo.

---

**Extended Data Figure 1:**
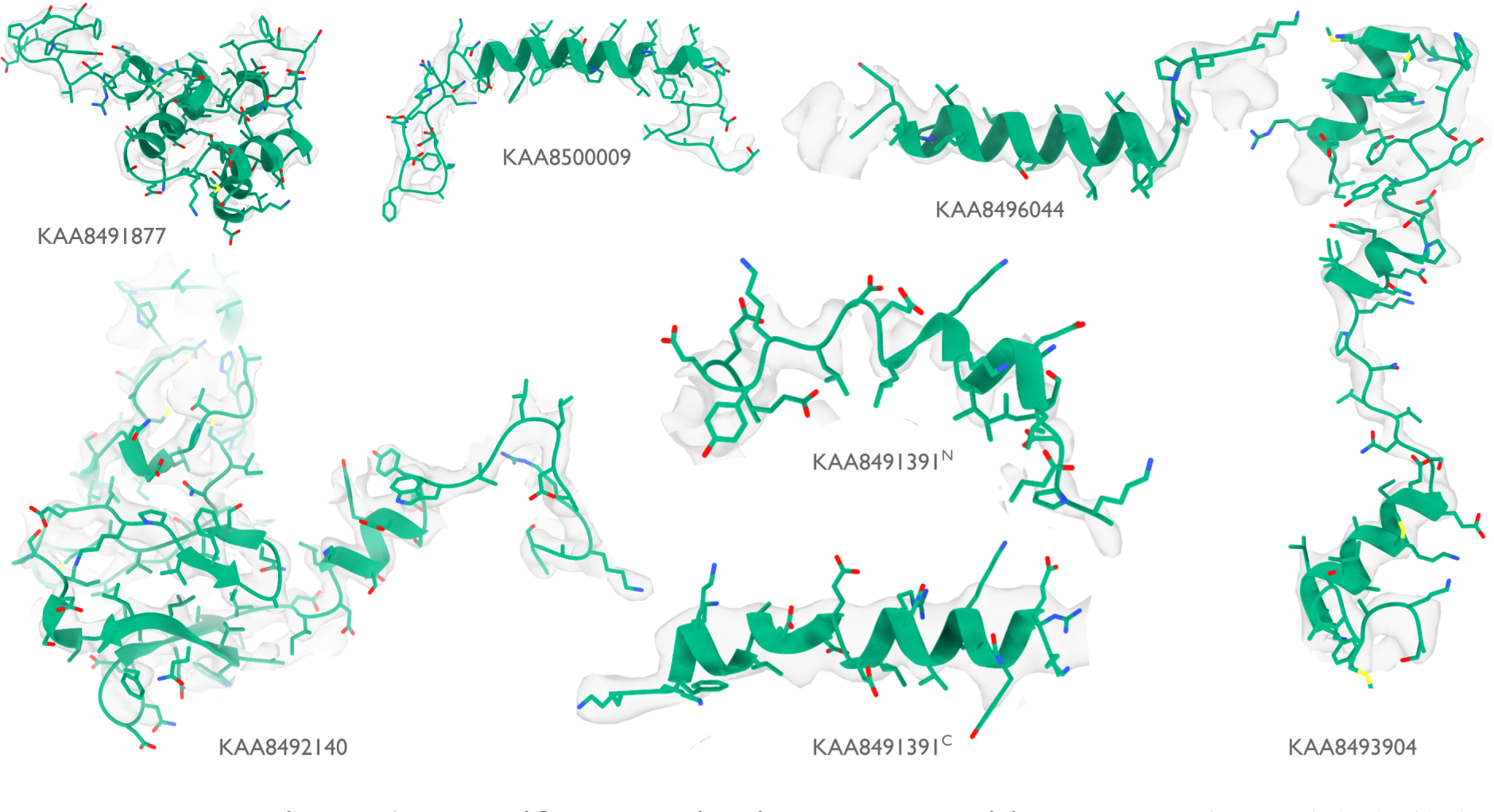
Identified proteins in the phycobilisome. Atomic models built by ModelAngelo (green) for the six proteins that were identified by ModelAngelo. Side chain densities in the cryo-EM map (transparent grey) are in agreement with those of the atomic models.

**Extended Data Figure 2:**
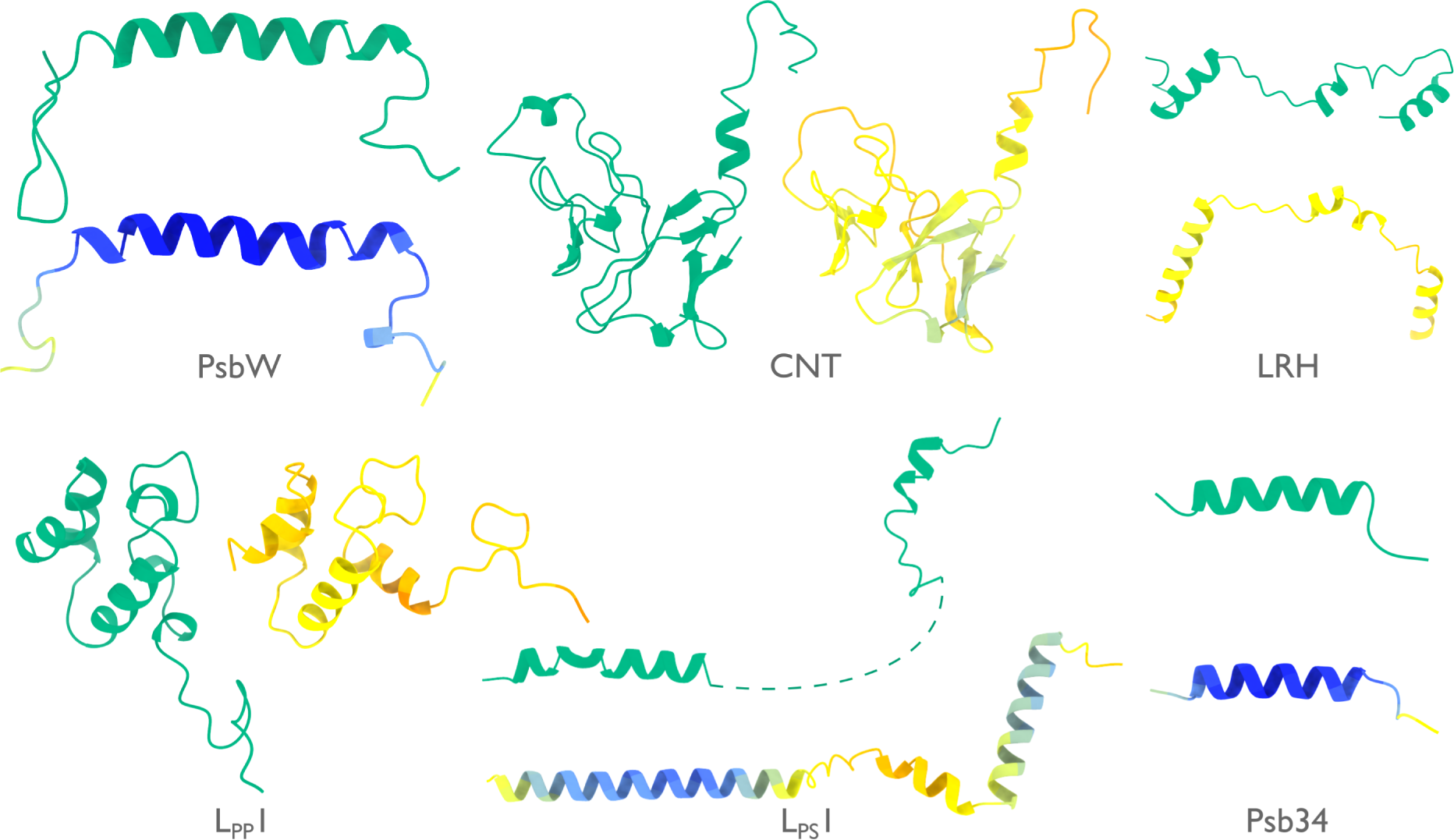
Models by ModelAngelo and AlphaFold for identified proteins in the phycobilisome. Models built by ModelAngelo (green) are shown next to predictions of the corresponding sequences by AlphaFold (*15*) (coloured by AlphaFold’s confidence from high in blue, to low in red).

**Extended Data Figure 3:**
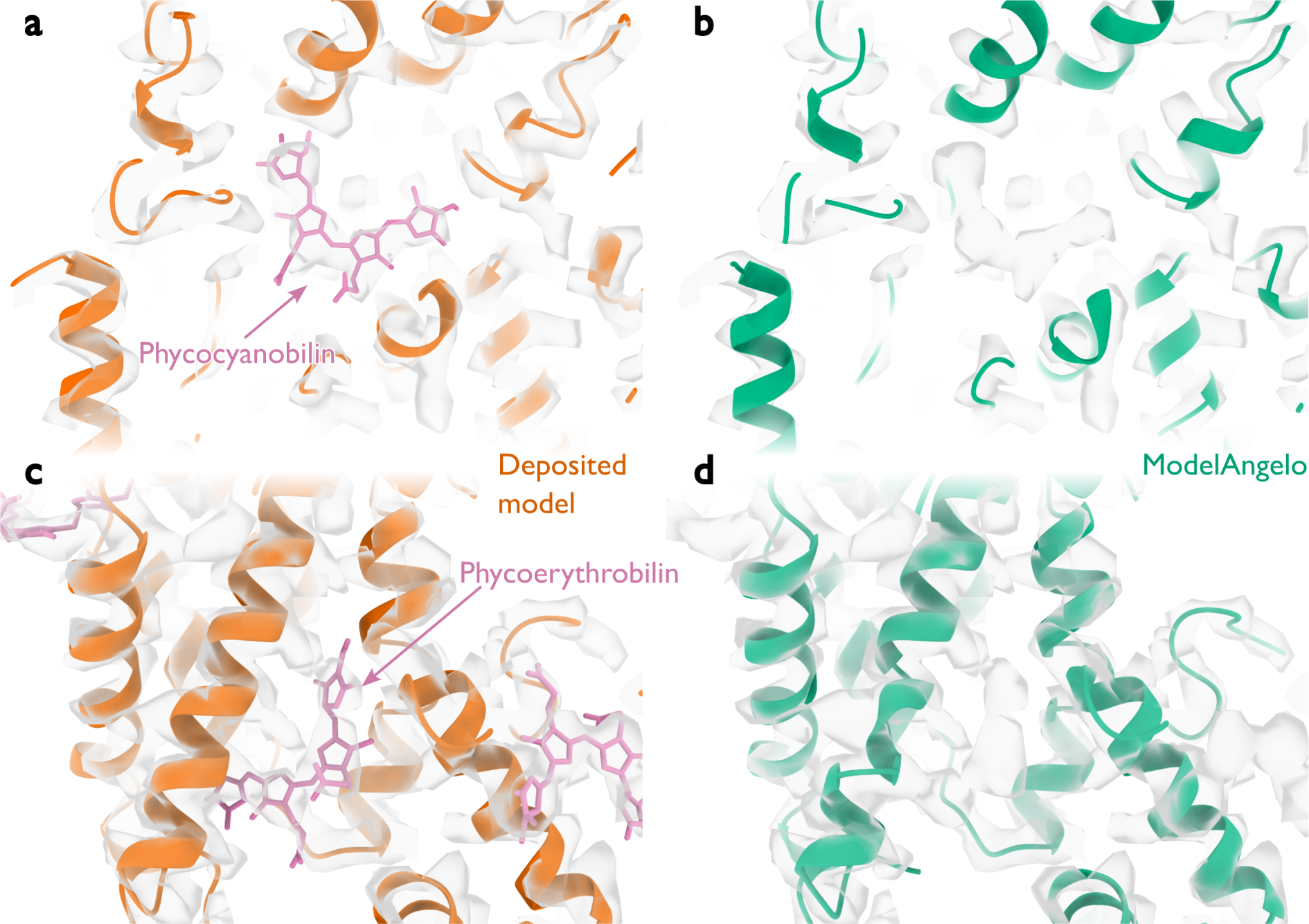
Performance around cofactors in the phycobilisome. **a**, Cartoon representation of protein backbones (orange) and stick representation of a phycocyanobilin cofactor (pink) in the cryo-EM density (transparent grey) for the deposited phycobilisome structure. **b**, as in panel a, but for the model built by ModelAngelo (green). ModelAngelo leaves the cofactor density empty. **c**, **d**, as in panels a, b but for a phycoerythrobilin cofactor.

**Extended Data Figure 4:**
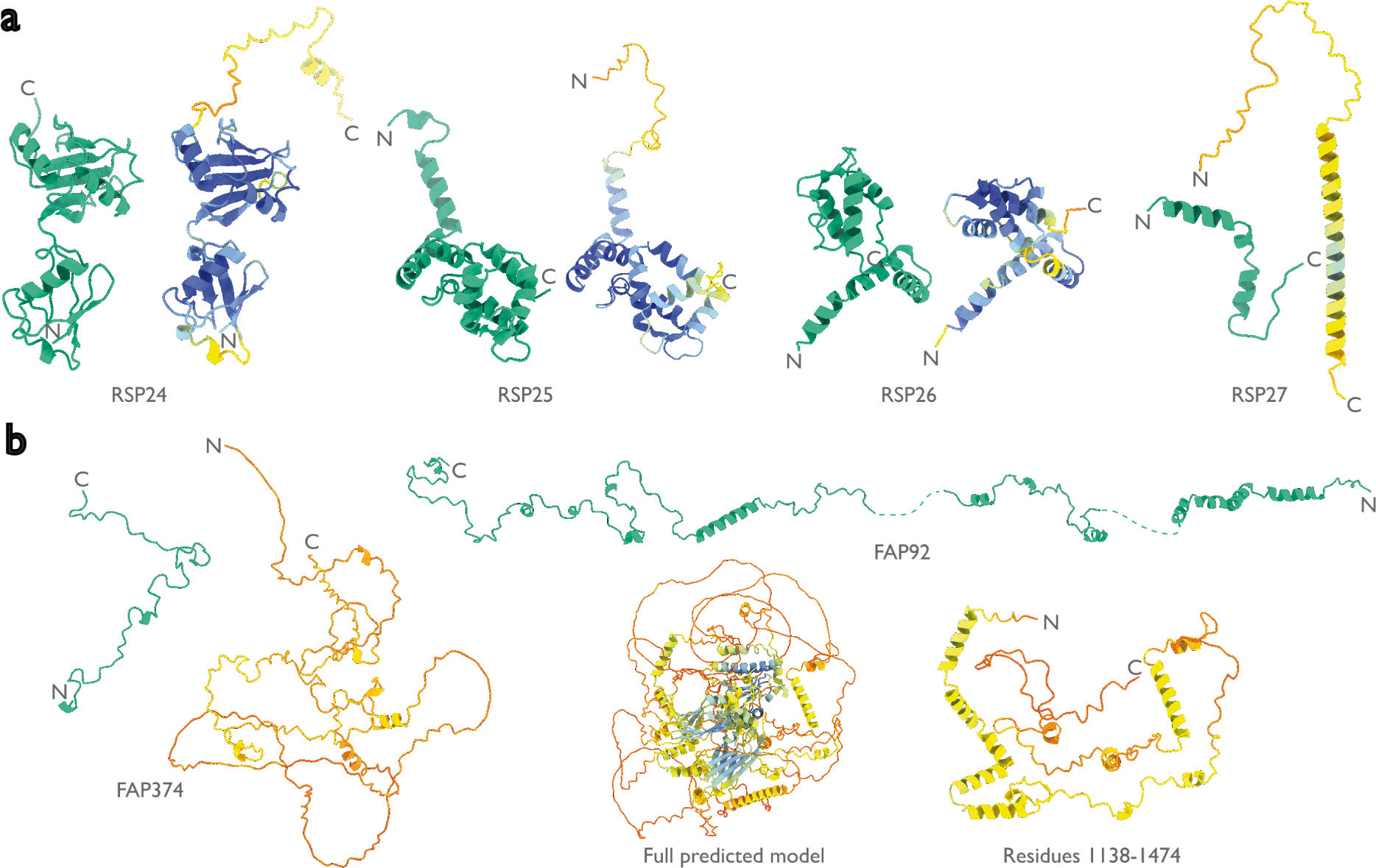
Models by ModelAngelo and AlphaFold for identified proteins in the ciliary axoneme. Models built by ModelAngelo (green) are shown next to predictions of the corresponding sequences by AlphaFold (*15*) (coloured by AlphaFold’s confidence from high in blue, to low in red). These are split between **a**, the radial spoke proteins, and **b**, the central apparatus microtubule proteins.

**Extended Data Figure 5:**
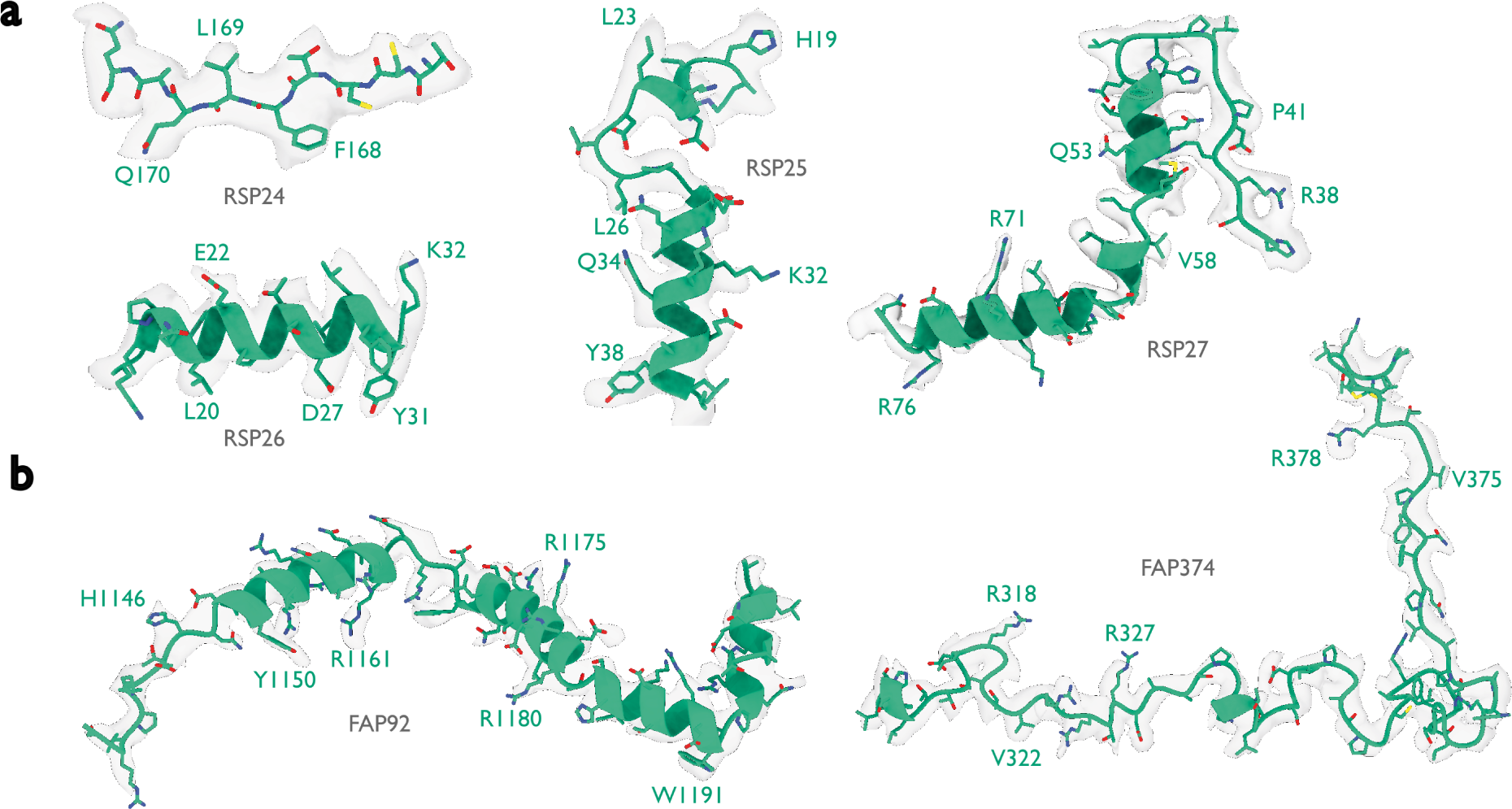
Identified proteins in the ciliary axoneme. Atomic models built by ModelAngelo (green) for the six proteins that were identified by ModelAngelo. Side chain densities in the cryo-EM map (transparent grey) are in agreement with those of the atomic models. These are split between **a**, the radial spoke proteins, and **b**, the central apparatus microtubule proteins.

**Extended Data Table 1:** Comparison with alternative approaches for the automated building of proteins. *MA* stands for ModelAngelo and *DT* for DeepTracer. *Calpha RMSD* is the root mean squared deviation of the predicted CA atoms against that of the deposition. *Backbone RMSD* is similar but for the CA, C, O and N atoms of the protein backbones. *Backbone recall* is the fraction of the deposited residues that were predicted to be within 3 Å (as measured between CA atoms). *Backbone precision* is the fraction of the predicted residues that have a corresponding residue present in the deposition within 3 Å . *Amino acid accuracy* is the fraction of the predicted residues that have a correctly predicted amino acid identity. Finally, *completeness* is the fraction of deposited residues that were predicted with the correct base annotation. Numbers indicated in boldface are the best in each metric.

| PDB | Res.<br>(Å) | Calpha<br>RMSD (Å) | Backbone<br>RMSD (Å) | Backbone<br>Recall | Backbone<br>Precision | Amino acid<br>Accuracy | Complete-<br>ness |
| --- | --- | --- | --- | --- | --- | --- | --- |
| 7uzs | 2.2 | **MA: 0.09**<br>DT: 0.37 | **MA: 0.11**<br>DT: 0.51 | MA: 0.92<br>**DT: 0.97** | MA: 0.96<br>**DT: 0.97** | **MA: 1.00**<br>DT: 0.99 | MA: 0.92<br>**DT: 0.96** |
| 7v0q | 2.5 | **MA: 0.09**<br>DT: 0.39 | **MA: 0.10**<br>DT: 0.64 | **MA: 0.99**<br>DT: 0.96 | **MA: 0.98**<br>DT: 0.96 | **MA: 1.00**<br>DT: 0.93 | **MA: 0.99**<br>DT: 0.89 |
| 7xgr | 2.6 | **MA: 0.30**<br>DT: 0.52 | **MA: 0.33**<br>DT: 0.83 | MA: 0.93<br>**DT: 0.94** | MA: 0.68<br>**DT: 0.94** | **MA: 0.99**<br>DT: 0.86 | **MA: 0.92**<br>DT: 0.81 |
| 7xjp | 2.71 | **MA: 0.31**<br>DT: 0.59 | **MA: 0.33**<br>DT: 1.13 | MA: 0.93<br>**DT: 0.94** | **MA: 1.00**<br>DT: 0.94 | **MA: 0.98**<br>DT: 0.59 | **MA: 0.91**<br>DT: 0.56 |
| 7xk4 | 3.1 | **MA: 0.19**<br>DT: 0.50 | **MA: 0.23**<br>DT: 0.71 | MA: 0.96<br>**DT: 0.97** | **MA: 1.00**<br>DT: 0.97 | **MA: 0.99**<br>DT: 0.86 | **MA: 0.95**<br>DT: 0.84 |
| 7xmv | 2.6 | **MA: 0.12**<br>DT: 0.35 | **MA: 0.15**<br>DT: 0.50 | **MA: 0.99**<br>DT: 0.98 | **MA: 1.00**<br>DT: 0.98 | **MA: 1.00**<br>DT: 0.97 | **MA: 0.99**<br>DT: 0.95 |
| 7xnz | 3.6 | **MA: 0.37**<br>DT: 0.65 | **MA: 0.42**<br>DT: 0.85 | **MA: 0.97**<br>DT: 0.96 | **MA: 0.99**<br>DT: 0.96 | **MA: 0.97**<br>DT: 0.89 | **MA: 0.94**<br>DT: 0.85 |
| 7yim | 2.6 | **MA: 0.41**<br>DT: 0.91 | **MA: 0.45**<br>DT: 1.48 | MA: 0.50<br>**DT: 0.70** | **MA: 0.98**<br>DT: 0.70 | **MA: 0.92**<br>DT: 0.14 | **MA: 0.46**<br>DT: 0.10 |
| 7ypx | 3.12 | **MA: 0.59**<br>DT: 0.84 | **MA: 0.67**<br>DT: 1.21 | **MA: 0.73**<br>DT: 0.58 | **MA: 0.97**<br>DT: 0.58 | **MA: 0.95**<br>DT: 0.56 | **MA: 0.69**<br>DT: 0.32 |
| 7zh0 | 3.2 | **MA: 0.45**<br>DT: 0.78 | **MA: 0.51**<br>DT: 1.36 | MA: 0.80<br>**DT: 0.93** | **MA: 0.98**<br>DT: 0.93 | **MA: 0.95**<br>DT: 0.66 | **MA: 0.77**<br>DT: 0.61 |
| 7zh6 | 3.67 | **MA: 0.55**<br>DT: 1.10 | **MA: 0.58**<br>DT: 1.41 | MA: 0.68<br>**DT: 0.91** | **MA: 0.98**<br>DT: 0.91 | **MA: 0.95**<br>DT: 0.41 | **MA: 0.65**<br>DT: 0.37 |
| 8a04 | 3.2 | **MA: 0.20**<br>DT: 0.45 | **MA: 0.25**<br>DT: 0.84 | **MA: 1.00**<br>DT: 0.95 | **MA: 1.00**<br>DT: 0.95 | **MA: 1.00**<br>DT: 0.92 | **MA: 1.00**<br>DT: 0.88 |
| 8a7d | 3.06 | **MA: 0.36**<br>DT: 0.73 | **MA: 0.42**<br>DT: 1.08 | MA: 0.79<br>**DT: 0.83** | **MA: 0.99**<br>DT: 0.83 | **MA: 0.97**<br>DT: 0.72 | **MA: 0.77**<br>DT: 0.59 |
| 8ap7 | 2.7 | **MA: 0.10**<br>DT: 0.33 | **MA: 0.12**<br>DT: 0.66 | MA: 0.97<br>**DT: 0.98** | MA: 0.91<br>**DT: 0.98** | **MA: 1.00**<br>DT: 0.97 | **MA: 0.97**<br>DT: 0.95 |
| 8ap8 | 3.7 | **MA: 0.31**<br>DT: 0.58 | **MA: 0.36**<br>DT: 0.86 | MA: 0.95<br>**DT: 0.97** | **MA: 0.99**<br>DT: 0.97 | **MA: 0.99**<br>DT: 0.79 | **MA: 0.94**<br>DT: 0.77 |
| 8avx | 3.5 | **MA: 0.68**<br>DT: 1.40 | **MA: 0.75**<br>DT: 2.02 | MA: 0.20<br>**DT: 0.81** | **MA: 0.99**<br>DT: 0.81 | **MA: 0.83**<br>DT: 0.09 | **MA: 0.16**<br>DT: 0.08 |
| 8bc2 | 2.6 | **MA: 0.13**<br>DT: 0.35 | **MA: 0.13**<br>DT: 0.50 | **MA: 1.00**<br>DT: 1.00 | **MA: 1.00**<br>DT: 1.00 | **MA: 1.00**<br>DT: 0.99 | **MA: 1.00**<br>DT: 0.98 |
| 8csw | 2.5 | **MA: 0.10**<br>DT: 0.39 | **MA: 0.11**<br>DT: 0.53 | **MA: 0.99**<br>DT: 0.95 | **MA: 0.97**<br>DT: 0.95 | **MA: 1.00**<br>DT: 0.93 | **MA: 0.99**<br>DT: 0.89 |
| 8cvz | 3.52 | **MA: 0.38**<br>DT: 0.80 | **MA: 0.43**<br>DT: 1.31 | MA: 0.78<br>**DT: 0.87** | **MA: 0.99**<br>DT: 0.87 | **MA: 0.96**<br>DT: 0.55 | **MA: 0.75**<br>DT: 0.47 |
| 8dh7 | 2.99 | **MA: 0.17**<br>DT: 0.47 | **MA: 0.19**<br>DT: 0.66 | **MA: 0.99**<br>DT: 0.99 | **MA: 1.00**<br>DT: 0.99 | **MA: 1.00**<br>DT: 0.87 | **MA: 0.99**<br>DT: 0.85 |
| 8dnm | 2.76 | **MA: 0.14**<br>DT: 0.44 | **MA: 0.17**<br>DT: 0.74 | **MA: 1.00**<br>DT: 0.99 | **MA: 1.00**<br>DT: 0.99 | **MA: 1.00**<br>DT: 0.97 | **MA: 1.00**<br>DT: 0.96 |
| 8dwi | 3.4 | **MA: 0.57**<br>DT: 0.83 | **MA: 0.61**<br>DT: 1.11 | **MA: 0.96**<br>DT: 0.96 | **MA: 0.96**<br>DT: 0.96 | **MA: 0.94**<br>DT: 0.56 | **MA: 0.90**<br>DT: 0.54 |
| 8dwu | 3.4 | **MA: 0.56**<br>DT: 0.95 | **MA: 0.62**<br>DT: 1.73 | MA: 0.35<br>**DT: 0.39** | **MA: 0.99**<br>DT: 0.39 | **MA: 0.94**<br>DT: 0.45 | **MA: 0.33**<br>DT: 0.18 |
| 8e50 | 3.67 | **MA: 0.34**<br>DT: 0.80 | **MA: 0.40**<br>DT: 1.21 | MA: 0.93<br>**DT: 0.96** | **MA: 1.00**<br>DT: 0.96 | **MA: 0.98**<br>DT: 0.49 | **MA: 0.91**<br>DT: 0.47 |
| 8efe | 3.8 | **MA: 0.68**<br>DT: 0.97 | **MA: 0.77**<br>DT: 1.57 | MA: 0.34<br>**DT: 0.66** | **MA: 0.99**<br>DT: 0.66 | **MA: 0.91**<br>DT: 0.21 | **MA: 0.31**<br>DT: 0.14 |
| 8evu | 2.58 | **MA: 0.10**<br>DT: 0.54 | **MA: 0.12**<br>DT: 0.84 | **MA: 1.00**<br>DT: 0.98 | **MA: 0.99**<br>DT: 0.98 | **MA: 1.00**<br>DT: 0.89 | **MA: 1.00**<br>DT: 0.87 |
| 8fma | 3.1 | **MA: 0.54**<br>DT: 2.16 | **MA: 0.58**<br>DT: 3.00 | **MA: 0.65**<br>DT: 0.21 | **MA: 0.97**<br>DT: 0.21 | **MA: 0.95**<br>DT: 0.06 | **MA: 0.62**<br>DT: 0.01 |

**Extended Data Table 2:** Comparison with alternative approaches for the automated building of nucleotides. *MA* stands for ModelAngelo, *CR* for CryoREAD, and *DT* for DeepTracer. *Phosphor RMSD* is the root mean squared deviation of the predicted P atoms against that of the deposition. *Backbone RMSD* is similar but for the OP1, P, OP2, and O5’ atoms of the nucleotide backbones. *Backbone recall* is the fraction of the deposited residues that were predicted to be within 3 Å (as measured between P atoms). *Backbone precision* is the fraction of predicted residues that have a corresponding residue present in the deposition within 3 Å . *Base accuracy* is the fraction of the predicted residues that have a correctly predicted nucleotide base. Finally, *completeness* is the fraction of deposited residues that were predicted with the correct base annotation. Numbers indicated in boldface are the best in each metric. Protofilaments 7-9

| PDB | Res.<br>(Å) | Phosphor<br>RMSD (Å) | Backbone<br>RMSD (Å) | Backbone<br>Recall | Backbone<br>Precision | Base<br>Accuracy | Complete-<br>ness |
| --- | --- | --- | --- | --- | --- | --- | --- |
| 7S1G | 2.48 | **MA: 0.36** | **MA: 0.48** | **MA: 0.96** | **MA: 0.99** | **MA: 0.80** | **MA: 0.77** |
|  |  | CR: 1.00 | CR: 1.99 | CR: 0.68 | CR: 0.66 | CR: 0.55 | CR: 0.37 |
|  |  | DT: 0.51 | DT: N/A | DT: 0.86 | DT: 0.56 | DT: N/A | DT: N/A |
| 7ZJX | 3.1 | **MA: 0.48** | **MA: 0.61** | **MA: 0.86** | **MA: 0.94** | **MA: 0.66** | **MA: 0.56** |
|  |  | CR: 1.24 | CR: 2.10 | CR: 0.72 | CR: 0.60 | CR: 0.53 | CA: 0.38 |
|  |  | DT: 0.86 | DT: N/A | DT: 0.61 | DT: 0.38 | DT: N/A | DT: N/A |
| 7ZPQ | 3.47 | **MA: 0.42** | **MA: 0.57** | **MA: 0.92** | **MA: 0.98** | **MA: 0.62** | **MA: 0.57** |
|  |  | CR: 1.14 | CR: 2.05 | CR: 0.72 | CR: 0.63 | CR: 0.52 | CR: 0.38 |
|  |  | DT: 0.56 | DT: N/A | DT: 0.76 | DT: 0.42 | DT: N/A | DT: N/A |

**Extended Data Table 3:** Proteins identified in the *C. reinhardtii* axoneme using ModelAngelo. X*^∗^*RSP25 and RSP26 are also expected to occur in the neck of RS2, which is thought to be identical to the neck of RS1 (*12*).

| Protein | Phytozome ID | Nr. of residues | Built residues | EMDB entry | Map res. (Å) | Location |
| --- | --- | --- | --- | --- | --- | --- |
| RSP24 | Cre08.g800895 | 226 | 1-187 | 22481 | 3.4 | RS2 stalk |
| RSP25 | Cre01.g034550 | 176 | 18-176 | 22475 | 3.2 | RS1 neck* |
| RSP26 | Cre17.g802036 | 128 | 2-125 | 22475 | 3.2 | RS1 neck* |
| RSP27 | Cre05.g240450 | 91 | 36-78 | 22475 | 3.2 | RS1 stalk |
| FAP92 | Cre13.g562250 | 1471 | 1138-1471 | 25381 | 3.8 | C1 microtubule/<br>Protofilaments 3-4 |
| FAP374 | Cre03.g176600 | 400 | 308-386 | 25381 | 3.8 | C1 microtubule/<br>Protofilaments 7-9 |

